# Lowland ecotype *Cyperus rotundus* L. affects growth and yield of rice under flooded conditions in the Philippines

**DOI:** 10.1101/2021.09.22.461350

**Authors:** Dindo King M. Donayre, Jessica Joyce L. Jimenez, Anna Maria Lourdes S. Latonio, Edwin C. Martin, Bhagirath S. Chauhan

## Abstract

Lowland ecotype *Cyperus rotundus* L. has been reported infesting irrigated lowland ricefields in the Philippines. Its effects on growth and yield of rice under flooded conditions are unknown. Two experimental runs were conducted in a screenhouse to determine the growth of lowland ecotype *C. rotundus* with transplanted rice and wet direct-seeded rice at a sowing density of 0, 22, 44, 66, and 88 tubers m^-2^; and its effect on growth and yield of rice. Except for height, growth variables of lowland ecotype *C. rotundus* were increased with the increase in its initial tuber densities. Compared with 22 tubers m^-2^, its number of off-shoots and tubers, and shoot and root biomass at 44 to 88 tubers m^-2^ increased by up to 3x. Growth variables of rice were reduced by the weed. Grain yield of transplanted rice was reduced by 14 to 38%; wet direct-seeded rice by 11 to 28%. Results suggest that lowland ecotype *C. rotundus* can grow well and reduce growth and yield of rice under flooded conditions. It also imply the need of developing a holistic weed control strategy against the weed.

## Introduction

Rice (*Oryza sativa* L.), ranked fourth after sugarcane (*Saccharum officinarum* L.), maize (*Zea mays* L.), and wheat (*Triticum aestivum* L.), is one of the most essential and most widely produced crops in the world [1]. In 2008, the estimated worldwide harvested area and production were 158,511,000 ha and 669,482,000 metric tons, which increased to 161,616,000 ha and 782,000,147 metric tons in 2018 [2]. These figures were estimated to increase 1% every year to meet the demand and feed the growing population of rice consumers.

Weeds are one of the omnipresent limiting factors to rice production [3]. Many farmers consider weeds as the most important pest particularly when it comes to the preparation and implementation of pest management strategies in the field [4] [5][4-8]. Uncontrollable growth of weeds can cause yield loss of rice by as much as 100% [9-12]. Thus, their heavy infestations in the field threaten the attainment of the potential yield of any cultivated rice.

*Cyperus rotundus* L. is a plant that belongs to the Cyperaceae family. It is a perennial and C_4_ plant that has horizontal and long-creeping rhizomes, slender thread-like stolons, and multiple -in-chained tubers [6] [7] [8][13-15]. It has been regarded as the world’s worst weed due to its reproductive behavior in the soil, flexibility for growing in many environmental conditions, and infestations in numerous crop production areas in the world [14,16]. Worldwide consolidated reports showed that it caused yield losses to many major crops by 6 to 100% [17-19]. In rice planted under upland conditions, full competition of the weed reduced grain yield by 43% under drill seeding and 51% under broadcast seeding [20]. In another study, *C. rotundus* at 750 plants m^-2^ reduced the grain yield of upland rice by 53% at 0 kg N ha^-1^, 44% at 600 kg N ha^-1^, and 47% at 120 kg N ha^-1^ [21].

*C. rotundus* has many ecotypes distributed around the world [22,23]. In the Philippines, earlier reports showed that it has a lowland ecotype that can grow in irrigated lowland ricefields [24-27]. Morphological characterization studies under laboratory and screenhouse conditions showed that it was taller, had more off-shoots, heavier biomass, and larger and heavier tubers as compared to its upland ecotype [24, 26, 28, 29]. Under field conditions, it was observed growing and infesting both transplanted and direct-seeded rice systems and was also taller (81.4 cm) than the cultivated rice plants (54.8 cm) [30]. It was elucidated that its survival under such conditions was due to its morphological and physiological adaptive mechanisms [28,29].

Knowledge of weed biology and ecology is one of the most important pre-requisites to selection and implementation of effective and economical weed management strategies. Information on growth of upland ecotypes *C. rotundus* and its negative effects on the growth and yield of rice were well-documented in previous studies under saturated conditions [20, 21]. For lowland ecotypes *C. rotundus*, however, such information is still lacking. Thus, this study was conducted to determine the growth of lowland ecotype *C. rotundus* in mixed culture with rice, and the weed effects on the growth and yield of transplanted and wet direct-seeded rice under flooded conditions (a simulation of irrigated lowland conditions).

## Materials and methods

Two experimental runs were conducted at the Philippine Rice Research Institute Central Experiment Station (PhilRice CES), Maligaya, Science City of Muñoz, Nueva Ecija, Philippines (N 15° 40’ 11.415”, E 120° 53’ 24.3802”). The first run (Run I) was conducted in February to May while the second run (Run II) was in August to November 2019. Since lowland ecotype *C. rotundus* was never been known to infest any experimental plots in PhilRice CES, the experiments then were conducted inside a screenhouse to easily monitor the growth and eradicate the weed afterwards. The screenhouse, where the experiment was established, was made of iron steel frames, a glass rooftop, and 1 cm^2^ steel mesh coverings on the sides. Weather conditions during the first run (Run I) had 27.5°C mean temperature (minimum = 22.8°C, maximum = 34.4°C), 75% relative humidity, 20.5 MJ m^-2^ solar radiation, and 9.2 h day^-1^ sunshine duration; the second run (Run II) had 27.1°C mean temperature (minimum = 23.8°C maximum = 32.6°C), 76% relative humidity, 14.4 MJ m^-2^ solar radiation, and 4.5 h day^-1^ sunshine duration.

### Preparation of rice plants

NSIC Rc 222, obtained from the Plant Breeding and Biotechnology Division of PhilRice CES, was used as the test rice variety due to its widespread usage among farmers across irrigated and rainfed-lowland areas in the country [31]. Five-hundred seeds were placed inside a fined-plastic net and soaked in a plastic container filled with clean tap water for 24 h. The seeds were pulled out from the water and incubated in the screenhouse for 72 h under cool conditions. Seeds with emerged radicles were seeded into a rectangular plastic container (length = 48 cm, width = 34 cm, depth = 15 cm) filled with 5 kg of moistened sterilized soil. The seeds were allowed to grow for 21 days under screenhouse conditions. Healthy, fully grown seedlings were selected and transplanted (referred as transplanted rice). In direct-seeding (referred as wet direct-seeded rice), seeds with emerged radicles were immediately seeded in each experimental unit.

### Preparation of *Cyperus rotundus*

Tubers of lowland ecotype *C. rotundus* were collected from irrigated-lowland ricefields of La Purisima, Aliaga, Nueva Ecija (15° 30’ 59” N, 12° 49’ 44” E). The area was selected based on the earlier reports [30]. Lowland ecotype *C. rotundus* plants, at the reproductive phase, were carefully pulled out from the soil using a shovel. Matured tubers were separated from the mother plants using scissors and placed inside a cylindrical plastic container. Collected tubers were then brought to the Weed Science Laboratory of the Crop Protection Division of PhilRice CES for washing and cleaning.

All cleaned tubers were placed in moistened autoclavable plastics (30 cm x 20 cm LW). The plastics were then placed in a temperature-controlled oven (Yamato DN83 constant temperature) for exposure of the tubers at 40°C for 4 h. The tubers were next transferred into a covered and moistened 50 L capacity storage plastic box for 5-day incubation at 25°C inside the laboratory. Tubers that sprouted (produced new shoots) were selected and planted in cylindrical plastic containers (30 cm diameter x 25 cm depth) previously filled with 5 kg sterilized moist soil. The tubers were allowed to grow and multiply into new plants under screenhouse conditions. After two months, matured tubers were harvested, cleaned, and kept in plastic containers. Newly harvested tubers were heat-exposed, incubated, and allowed to sprout again under the same process as described above. Tubers that sprouted were selected; grouped into large (*x*^-^ =26.1 mm × 11.1 mm LW), medium (*x*^-^ =16.7 mm × 9.7 mm LW), and small (*x*^-^ =11.6 mm × 8.5 mm LW); and planted in each experimental unit. Grouping was done to allow each treatment to receive all the sizes of tubers during planting.

### Experimental treatments and design

An additive design, in which the density of the crop was kept constant while that of the weed was increased, was utilized in this experiment to determine the outcome of rice - lowland ecotype *C. rotundus* competition [32]. The experimental unit used was a black, 0.23 m^2^ rectangular plastic box (length = 0.54 m, width = 0.42 m, and deep = 0.36 m) filled with 62 kg moist soil (Maligaya soil series, 0.104 N%, 20.1 ppm P, 0.16 me K/100 g soil, pH=5.5). Each box was either transplanted with eight seedlings (21-day-old) at 20 cm x 20 cm distance or seeded with 36 pre-germinated rice seeds (3-day-old) at 8 cm x 8 cm distance. Sprouted tubers of lowland ecotype *C. rotundus* were then simultaneously planted in boxes at 0, 5, 10, 15, and 20 tubers which were equivalent to 22, 44, 66, and 88 tubers m^-2^. Lowland ecotype *C. rotundus* at 5, 10, 15, and 20 tubers were planted at distances of 20 cm x 20 cm, 15 cm x 15 cm, 12 cm x 12 cm, and 10 cm x 10 cm, respectively. At 7 days after planting (DAP), 33 L of clean tap water was supplied in each box and maintained thereafter at 5 cm depth. Rice and lowland ecotype *C. rotundus* plants were allowed to grow until the crop’s maturity stage. All plants in each box were nourished with fertilizers at three periods of application (15, 30, and 45 DAP) following the recommended rate of 120-40-40 nitrogen, phosphorus, and potassium. Boxes with either transplanted or direct-seeded rice and sprouted tubers of lowland ecotype *C. rotundus* at different densities were separately arranged in a randomized complete block design with five replications [33].

### Measurements

Height of lowland ecotype *C. rotundus* was measured using a meter stick while the numbers of off-shoots were manually counted at 60 days after transplanting or direct-seeding of rice. After harvesting, dry shoot and root biomass box^-1^ were determined by drying the fresh biomass inside an oven (Yamato DN83 constant temperature oven) for 48 h at 70°C and weighing the dried biomass using a digital weighing balance (A&D Electronic Balance FX-3000). Number of tubers box^-1^ was also determined by sieving the soil in each box and then collecting and manually counting the tubers. All collected data were converted into per square meter (m^-2^). Fold increases (times) on growth variables of the weed were then calculated using the formulae below:

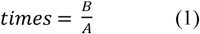

where B served as the mean values at 44, 66, and 88 tubers m^-2^ and A as the mean values at 22 tubers m^-2^.

Height, number of tillers and leaves plant^-1^ of rice were also manually measured and counted at 60 days after transplanting or direct-seeding. Dried shoot and root biomasses box^-1^ were also recorded following the same procedure as that of lowland ecotype *C. rotundus*. Yield components such as number of panicles, number of grains per panicle, percentage filled spikelets, and 1000-grain weight were also collected after harvest. Grain yield was then calculated from yield components and adjusted to 14% moisture content [34]. All collected data were also converted into per square meter (m^-2^). Reductions on growth and yield variables of rice were calculated using the equation below [35]:

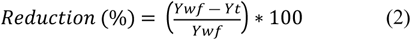

where Y_wf_ as mean values on growth and yield variables of rice at 0 tuber, and Y_t_ as mean values on growth and yield variables of rice at 22, 44, 66, and 88 tubers m^-2^, respectively.

### Statistical analysis

Data on the growth of the weed as well as growth and yield of rice in two runs were subjected to combined ANOVA using the Statistical Tool for Agricultural Research (STAR 201) of the International Rice Research Instititute. All means were then compared using the least significant difference (LSD) test at the 5% level of significance. A two-sample t-test was also utilized to compare the fold increase of lowland ecotype *C. rotundus*, and the grain yields and yield losses between transplanted rice and DSR at each level of initial tuber density. A Pearson-product moment correlation coefficient was also computed to determine the strengths and direction of relationships of each growth variable of lowland ecotype *C. rotundus*; and growth variables of lowland ecotype *C. rotundus* on grain yield of both transplanted rice and wet direct-seeded rice. To determine if the tuber density, number of off-shoots, and shoot biomass m^-2^ of the weed were significant contributors and predictors to grain yield of both transplanted and wet direct-seeded rice, values of each variable were fitted to a second-order polynomial quadratic equation using the Analysis ToolPak of Microsoft Excel 2016. The equation was calculated as shown below [36]:

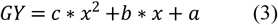

where GY is the grain yield of transplanted /wet direct-seeded rice, *a* is the intercept on the y-axis, *b* is the slope of the curve, *c* is the quadratic term of the model, and x is growth variables of the weed.

To estimate grain yield losses due to lowland ecotype *C. rotundus*, data on grain yield loss of transplanted and wet direct-seeded rice in response to tuber density, number of off-shoots, and shoot biomass m^-2^ of lowland ecotype *C. rotundus* were fitted to a rectangular hyperbola equation using the Mycurvefit computer program (Mycurvefit.com). The three growth variables of the weed were chosen because they are easy to observe, collect, and apply in times of practicality and immediate predictions in the field. The equation of rectangular hyperbola was calculated as shown below [37]:

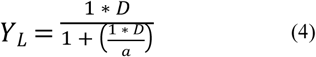

where Y_L_ as the grain yield loss of transplanted or wet direct-seeded rice; *D* as the tuber density, number of off-shoots, and shoot biomass; *i* as the percentage yield loss per weed plant per unit area as initial tuber density, number of off-shoots, and shoot biomass approaches zero; and *a* as the percentage yield loss as tuber density, number of off-shoots, and shoot biomass approaches infinity. All means of the respective growth variables then fitted against the values of respective grain yield reductions.

## Results

### Growth of lowland ecotype *Cyperus rotundus* with rice

The interaction effect of run and density was significant on number of off-shoots (*P*<0.01), number of tubers (*P*<0.02), and shoot biomass (*P*<0.02) of lowland ecotype *C. rotundus* grown with transplanted rice. In wet direct-seeded rice, the interaction effect of run and density was not significant on all growth variables of the weed. The main effect of run was significant on number of off-shoots (*P*<0.001) and number of tubers (*P*<0.001) while the main effect of density was significant on all growth variables (all *P*<0.001) except for the height.

Height of lowland ecotype *C. rotundus* had no significant differences when grown with transplanted rice at 22, 44, 66, and 88 tubers m^-2^ (**Table 1**). But its number of off-shoots, number of tubers, and shoot biomass significantly differed. The three growth variables of the weed significantly increased as the number of tuber density m^-2^ increased. The increase, however, was much higher during the second run (Run II). In the first run (Run I), compared with 22 tubers m^-2^, the number of off-shoots increased by 1.7 to 3.2 times, no. of tubers by 1.5 to 2.8 times, and shoot biomass by 1.6 to 3 times at 44, 66, and 88 m^-2^, respectively. In Run II, the number of off-shoots increased by 2.1 to 3.3 times, no. of tubers by 2.2 to 3.1 times, and shoot biomass by 2.3 to 2.9 times. Root biomass increased by 1.7 to 2.6 times.

**Table 1.** Growth of lowland ecotype *Cyperus rotundus* in association with rice

| Tuber Density<br>$\text{m}^{-2}$ | Height<br>(cm plant<sup>-1</sup>) | Number of off-shoots plant<sup>-1</sup> | Number of<br>tubers plant<sup>-1</sup> | Shoot biomass<br>(g $\text{m}^{-2}$) | Root biomass<br>($\text{m}^{-2}$) |
| --- | --- | --- | --- | --- | --- |
| Transplanted Rice |  |  |  |  |  |
| 22 | 173<sup>a</sup> | 162<sup>c</sup> (219<sup>d</sup>) | 389<sup>d</sup> (374<sup>c</sup>) | 297<sup>c</sup> (380<sup>c</sup>) | 74<sup>d</sup> |
| 44 | 175<sup>a</sup> | 276<sup>b</sup> (465<sup>c</sup>) | 589<sup>c</sup> (825<sup>b</sup>) | 490<sup>b</sup> (890<sup>b</sup>) | 125<sup>c</sup> |
| 66 | 173<sup>a</sup> | 333<sup>b</sup> (584<sup>b</sup>) | 780<sup>b</sup> (1083<sup>a</sup>) | 581<sup>b</sup> (1043<sup>ab</sup>) | 157<sup>b</sup> |
| 88 | 169<sup>a</sup> | 521<sup>a</sup> (714<sup>a</sup>) | 1090<sup>a</sup> (1150<sup>a</sup>) | 898<sup>a</sup> (1092<sup>a</sup>) | 193<sup>a</sup> |
| LSD | - | 82 | 139 | 184 | 32 |
| Wet Direct-Seeded Rice |  |  |  |  |  |
| 22 | 163<sup>a</sup> | 148<sup>d</sup> | 301<sup>d</sup> | 220<sup>d</sup> | 56<sup>d</sup> |
| 44 | 167<sup>a</sup> | 270<sup>c</sup> | 569<sup>c</sup> | 404<sup>c</sup> | 89<sup>c</sup> |
| 66 | 163<sup>a</sup> | 363<sup>b</sup> | 752<sup>b</sup> | 583<sup>b</sup> | 124<sup>b</sup> |
| 88 | 164<sup>a</sup> | 435<sup>a</sup> | 945<sup>a</sup> | 663<sup>a</sup> | 144<sup>a</sup> |
| LSD | - | 43 | 117 | 63 | 19 |

Values from Run II (inside parenthesis) are shown due to interaction effects

Similarly, height of lowland ecotype *C. rotundus* had no significant differences when grown with wet direct-seeded rice at 22, 44, 66, and 88 tubers m^-2^. Its growth variables also increased with the increase in tuber density. Compared with 22 tubers m^-2^, the number of off-shoots at 44 to 88 tubers m^-2^ increased by 1.8 to 2.9 times, no. of tubers by 1.9 to 3.1 times, shoot biomass by 1.8 to 3 times, and root biomass by 1.6 to 2.6 times.

Results of statistical analysis using the two-sample t-test showed that growth variables of lowland ecotype *C. rotundus* with wet direct-seeded rice were significantly lower in terms of height at 44 tubers m^-2^ (*P*<0.05); number of off-shoots at 44 (*P*<0.05) and 88 tubers m^-2^ (*P*<0.001); number of tubers at 88 m^-2^ (*P*<0.05); shoot biomass at 22 (*P*<0.05), 44 (*P*<0.001), and 88 (*P*<0.001) tubers m^-2^; and root biomass at 44 (*P*<0.05) and 88 tubers m^-2^ (*P*<0.05). Its growth variables with wet direct-seeded rice, however, were not significantly different with transplanted rice when it comes to height at 22, 44, and 88 tubers m^-2^; number of off-shoots at 22 and 44 tubers m^-2^; number of tubers at 22, 44, and 66 tubers m^-2^; shoot biomass at 66 tubers m^-2^; and root biomass at 22 and 66 tubers m^-2^.

### Effects of lowland ecotype *Cyperus rotundus* on growth and yield of rice

The interaction effect of run and density was only significant on number of leaves (*P*<0.05), number of tillers (*P*<0.05), and number of panicles (*P*<0.05) of transplanted rice. The main effect of run was significant on height (*P*<0.001), shoot biomass (*P*<0.001), root biomass (*P*<0.001), number of grains panicle^-1^ (*P*<0.05), filled spikelets (*P*<0.001), and grain yield (*P*<0.001). The the main effect of density, on the other hand, was only significant on height (*P*<0.05), shoot biomass (*P*<0.001), root biomass (*P*<0.000) and grain yield (<0.001).

In wet direct-seeded rice, the interaction effect of run and density was only significant on number of tillers (*P*<0.05). The main effect of the run was significant on shoot and root biomasses (*P*<0.05, <0.05), number of panicles (*P*<0.001), number of grains panicle^-1^ (*P*<0.05), filled spikelets (*P*<0.05), and grain yield (*P*<0.05). The main effect of density was only significant on height (*P*<0.001), number of leaves (*P*<0.05), shoot biomass (*P*<0.001), root biomass (*P*<0.001), and grain yield (*P*<0.001).

Growth and yield variables of transplanted rice were highest when lowland ecotype *C. rotundus* was absent (0 tuber m^-2^) (**Table 2**). When the weed was present at 22, 44, 66, and 88 tubers m^-2^, its growth and yield were significantly reduced except for filled spikelets and 1000-grain weight. Among the variables, its number of leaves and tillers, shoot and root biomasses, number of panicles, and grain yield were the most highly affected; height and number of grains panicle^-1^ were the least. At 22, 44, 66, and 88 tubers m^-2^, its height was reduced by 2.5, 3.7, 2.0, and 5.5%; number of leaves by 3.5, 19.5, 30.7, and 33.1% in Run I while 40.1, 39.5, 43.3, and 54.4% in Run II; number of tillers by 0.3, 18.8, 29.4, and 31.7% in Run I and 36.2, 34.5, 41.1, and 51.6% in the Run II; shoot biomass by 22.6, 37.3, 41.8, and 54.2%; root biomass by 39.1, 48.9, 51.0, and 60.2%; number of panicles by 0, 18.7, 21.7, and 32.2% in Run I and 35.9, 40.5, 47.1, and 51.7% in Run I; number of grains panicle^-1^ by 7.5, 8.7, 6.7, and 14.5%; and grain yield by 13.5, 27.3, 27.9, and 38.2% (**Fig. 1**), respectively. Reductions in growth and yield were greatest at 88 tubers m^-2^.

**Table 2.** Growth and yield components of rice as affected by different densities of lowland ecotype *Cyperus rotundus* under flooded conditions

| Tuber Density m<sup>-2</sup> | Height (cm plant<sup>-1</sup>) | No. of leaves plant<sup>-1</sup> | No. of tillers plant<sup>-1</sup> | Shoot biomass (g m<sup>-2</sup>) | Root biomass (g m<sup>-2</sup>) | No. of panicles m<sup>-2</sup> | No. of grains panicle<sup>-1</sup> | Filled spikelets (%) | 1000 grain weight (g) |
| --- | --- | --- | --- | --- | --- | --- | --- | --- | --- |
| Transplanted Rice |  |  |  |  |  |  |  |  |  |
| 0 | 122<sup>a</sup> | 77<sup>a</sup> (82<sup>a</sup>) | 15<sup>a</sup> (14<sup>a</sup>) | 1103<sup>a</sup> | 352<sup>a</sup> | 405<sup>a</sup> (342<sup>a</sup>) | 203<sup>a</sup> | 48<sup>a</sup> | 22<sup>a</sup> |
| 22 | 119<sup>abc</sup> | 74<sup>a</sup> (49<sup>b</sup>) | 15<sup>a</sup> (9<sup>bc</sup>) | 854<sup>b</sup> | 215<sup>b</sup> | 408<sup>a</sup> (219<sup>b</sup>) | 188<sup>a</sup> | 51<sup>a</sup> | 23<sup>a</sup> |
| 44 | 118<sup>bc</sup> | 62<sup>b</sup> (49<sup>b</sup>) | 12<sup>b</sup> (9<sup>b</sup>) | 691<sup>bc</sup> | 180<sup>b</sup> | 330<sup>b</sup> (203<sup>bc</sup>) | 185<sup>a</sup> | 51<sup>a</sup> | 22<sup>a</sup> |
| 66 | 119<sup>ab</sup> | 53<sup>b</sup> (46<sup>bc</sup>) | 10<sup>b</sup> (8<sup>bc</sup>) | 642<sup>cd</sup> | 173<sup>b</sup> | 317<sup>b</sup> (181<sup>bc</sup>) | 189<sup>a</sup> | 53<sup>a</sup> | 22<sup>a</sup> |
| 88 | 115<sup>c</sup> | 51<sup>b</sup> (37<sup>c</sup>) | 10<sup>b</sup> (7<sup>c</sup>) | 505<sup>d</sup> | 140<sup>b</sup> | 275<sup>c</sup> (165<sup>c</sup>) | 173<sup>a</sup> | 54<sup>a</sup> | 23<sup>a</sup> |
| LSD | 4 | 11 | 2 | 165 | 81 | 39 | - | - | - |
| Wet Direct-Seeded Rice |  |  |  |  |  |  |  |  |  |
| 0 | 100<sup>a</sup> | 25<sup>a</sup> | 5<sup>a</sup> (4<sup>a</sup>) | 960<sup>a</sup> | 204<sup>a</sup> | 499<sup>a</sup> | 111<sup>a</sup> | 67<sup>a</sup> | 21<sup>a</sup> |
| 22 | 99<sup>ab</sup> | 24<sup>ab</sup> | 5<sup>b</sup> (4<sup>a</sup>) | 928<sup>a</sup> | 158<sup>ab</sup> | 447<sup>b</sup> | 115<sup>a</sup> | 64<sup>a</sup> | 21<sup>a</sup> |
| 44 | 97<sup>bc</sup> | 22<sup>bc</sup> | 4<sup>bc</sup> (3<sup>b</sup>) | 807<sup>b</sup> | 126<sup>bc</sup> | 417<sup>b</sup> | 112<sup>a</sup> | 64<sup>a</sup> | 22<sup>a</sup> |
| 66 | 95<sup>d</sup> | 21<sup>c</sup> | 4<sup>bc</sup> (4<sup>ab</sup>) | 734<sup>b</sup> | 121<sup>bc</sup> | 373<sup>c</sup> | 111<sup>a</sup> | 62<sup>a</sup> | 23<sup>a</sup> |
| 88 | 96<sup>cd</sup> | 20<sup>c</sup> | 4<sup>c</sup> (4<sup>ab</sup>) | 624<sup>c</sup> | 96<sup>c</sup> | 358<sup>c</sup> | 109<sup>a</sup> | 66<sup>a</sup> | 22<sup>a</sup> |
| LSD | 2 | 2 | 1 | 109 | 48 | 32 | - | - | - |

Values from Run II (inside parenthesis) are shown due to interaction effects

**Fig. 1.**
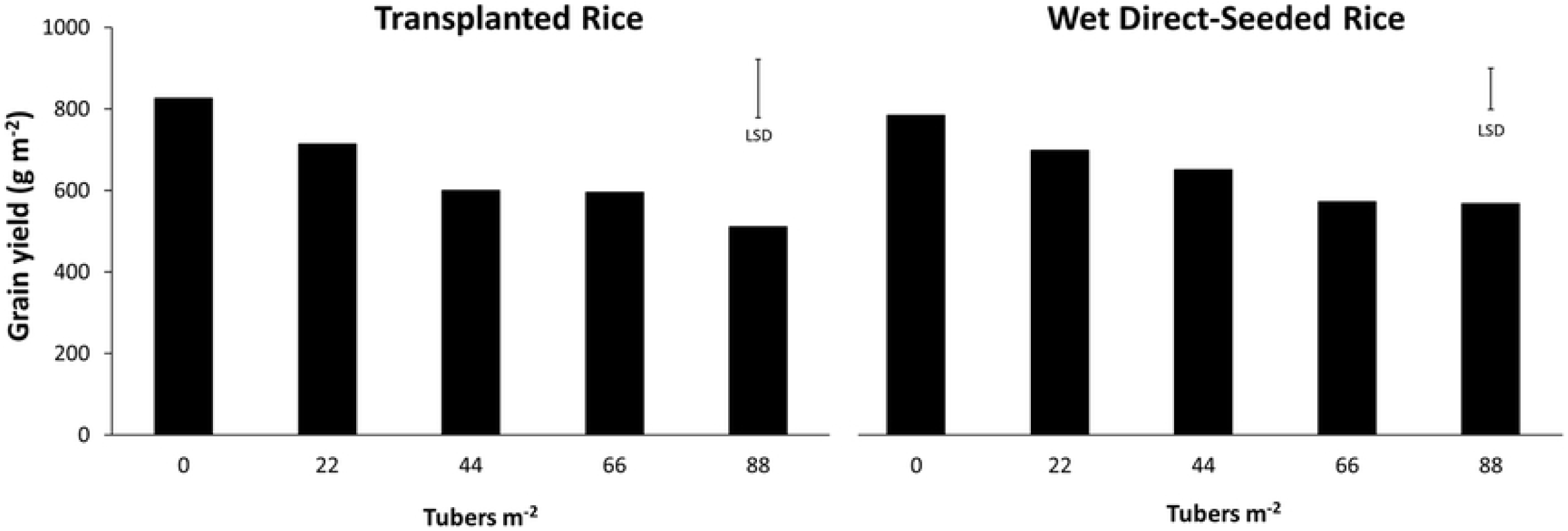
Grain yield of rice as affected by different densities of lowland ecotype *Cyperus rotundus* under flooded conditions

Growth and yield of wet direct-seeded rice were also highest when lowland ecotype *C. rotundus* was absent. When the weed was present, however, its number of leaves and tillers, shoot and root biomasses, number of panicles, and grain yield were then affected. Its height was less affected while its number of grains panicle^-1^, filled spikelets, and 1000-grain weight were not affected. At 22, 44, 66, and 88 tubers m^-2^, its height was reduced by 0.9, 2.6, 5.1, and 4.3%; number of leaves by 4.6, 10.1, 16, and 18.4%; number of tillers by 12.4, 16.6, 20.5, and 26.3% in Run I while 1, 16, 9.5, and 8.5% in Run II; shoot biomass by 3.3, 15.9, 23.6, and 35%;root biomass by 22.9, 38.3, 40.7, and 53.2%; number of panicles by 10.3, 16.3, 25.2, and 28.2%; and grain yield by 11, 17, 27, and 27.6%, respectively. Reductions in growth and yield were also greatest at 88 tubers m^-2^. Although the reductions on grain yields of wet direct-seeded rice were generally lower compared to transplanted rice, results of statistical analysis through TTEST showed no significant differences (data not shown).

### Relationships and yield loss estimations

Grain yields of transplanted rice and wet direct-seeded rice were strongly but negatively correlated to tuber densities (*R* = -0.968, -0.972), number of off-shoots (*R* = -0.991, -0.989), number of tubers (*R* = -0.986, -0.983), shoot biomass (*R* = -0.995, -0.994), and root biomass (*R*= -0.990, -0.993) of lowland ecotype *C. rotundus*. Height of the weed was moderately correlated to grain yield of both transplanted rice and wet direct-seeded rice (*R* = -0.795). Results of the analysis using polynomial quadratic regression also indicated that 96.9, 98.9, 97.5, 98.9, and 98% of grain yield of transplanted rice; and 98.3, 95.9, 97.8, 95.6, and 97% grain yield of wet direct-seeded rice were explained by the initial tuber densities, number of off-shoots, number of tubers, shoot biomass, and root biomass. Furthermore, the analysis also showed that growth variables of the weed were significant predictors and contributors to the grain yield of transplanted rice (*P* = 0.031, 0.011, 0.025, 0.011, and 0.020) as well as that of wet direct-seeded rice (*P* = 0.017, 0.020, 0.026, 0.010, and 0.013) at 5% level of significance.

Among the three growth variables of lowland ecotype *C. rotundus*, only tuber density and number of off-shoots fitted significantly in rectangular hyperbola (**Table 3**). The estimated yield losses of transplanted rice per weed were 0.8 and 0.1% as the tuber density and number of off-shoots of the weed approached zero (*i*); 75.7 and 66.1% as the two growth variables approached infinity (*a*). On the other hand, the estimated yield losses of wet direct-seeded rice per weed were 0.6 and 0.1% as the two growth variables approached *i* while 63.8 and 100% *a*, respectively.

**Table 3.** Rectangular hyperbola model parameter estimates for yield losses of rice in competition with lowland ecotype *Cyperus rotundus* under flooded conditions

| Rice | Parameter estimates (%) | | $R^2$ | $P$ |
| --- | --- | --- | --- | --- |
| | $i$ | $A$ | | |
| Transplanted Rice |  |  |  |  |
| Tuber density | 0.8 | 75.8 | 97.3 | 0.004** |
| No. of off-shoots | 0.1 | 66.1 | 87.5 | 0.001** |
| Shoot biomass (g) | 0.1 | 26.7 | 50.7 | 0.083<sup>ns</sup> |
| Wet Direct-seeded Rice |  |  |  |  |
| Tuber density | 0.6 | 63.8 | 97.9 | 0.003** |
| No. of off-shoots | 0.1 | 100 | 98.0 | 0.003** |
| Shoot biomass (g) | 0.1 | 100 | 98.0 | 0.022<sup>ns</sup> |

*i*- yield loss per weed plant per unit area as the initial tuber density, no. of off-shoots, and shoot biomass approached zero; *a*- yield loss as the initial tuber density, no. of off-shoots, and shoot biomass approached infinity; *R*<sup>2</sup>- coefficient of determination; *P* values at 5% level of significance, \*\**P*<.01 - highly significant, ns- not significant

## Discussion

Knowledge on weed biology and ecology helps understand the strengths and weaknesses of a certain weed. It also helps plan and execute effective weed management strategies. Our study demonstrated that height of lowland ecotype *C. rotundus* was not influenced by neither its increasing initial tuber density nor the presence of transplanted or wet direct-seeded rice, suggesting that light was not a limiting factor for intraspecific and interspecific competitions. In fact, our findings are similar to a interspecific competition study where height of upland *C. rotundus* was not affected at density ratios of 1:12 and 1:24 (*C. rotundus*:rice) [38]. In a intraspecific study, however, height of *C. rotundus* increased significantly as its density was increased from 1 plant pot^-1^ to 9 and 25 plants pot^-1^ [39].

Lowland ecotype *C. rotundus* grew and produced numerous off-shoots and tubers side by side with rice suggesting that it can indeed flourish and survive under flooded conditions. Our previous study on regrowth capability of lowland ecotype *C. rotundus* under flooded conditions also showed that 50 and 40% of its pre-sprouted tubers and 20% of its non-sprouted tubers abled to germinate and grow when planted at the soil surface (0 cm burial depth) and flooded with water at 3 to 5 cm depths [40]. No continuous growths, however, were observed on tubers subjected to flooding depths of 3 cm and 5 cm buried at 5 cm and 10 cm.

Previous study also had similar findings on *C. rotundus* that had been collected from rice-onion cropping systems in Nueva Ecija, Philippines [26]. It was reported that both of its lowland and upland types, buried at 1 cm depth, sprouted and emerged at all water levels (1, 3, and 5 cm). Between the two, the height and fresh weight of the lowland ecotype were less reduced at 5 cm flooding. It was elucidated that the tolerance of lowland ecotype *C. rotundus* to flooding was due to its large carbohydrate content, amylase activity, and ability to maintain high levels of soluble sugars in the tubers during germination and early growth [29]. These activities were enhanced with modulation of alcohol dehydrogenase and pyruvate decarboxylase to regulate utilization of carbohydrate reserves and sustain substrate supply to keep away from starvation and death of seedlings under prolonged flooding.

It was also elaborated that the adaptation of lowland ecotype *C. rotundus* in flooded conditions was due to its bigger tubers that have high carbohydrate and soluble sugar contents, larger stems, and more air spaces (aerenchyma) that enable oxygen diffusion to its submerged parts, capacity to mobilize and utilize carbohydrate reserves for energy production through anaerobic respiration, capacity to optimize use of carbohydrate reserves through regulation of key enzymes like the alcohol dehydrogenase; and down-regulation of lactate dehydrogenase possibly to prevent lactate accumulation and cellular acidosis [28].

It was reported that *C. rotundus* produced more shoots and tubers during the first run of an experiment that had high solar radiation (406 cal cm^2^) than during the second run that had low solar radiation (270 cal cm^2^) [39]. Our experiment, however, had opposing results and showed that lowland ecotype *C. rotundus*, grown with transplanted rice, produced more off-shoots and tubers as well as shoot biomass despite the occurrence of low solar radiation (Run II). This growth response of the weed could be viewed as a way of plasticity since low solar radiation would mean less entrapment of light for photosynthesis. Lowland ecotype *C. rotundus* has thin and fewer leaves, thus, it has to overcome these barriers by producing more off-shoots to have more leaves that will eventually trap more light energy for photosynthesis. More off-shoots then resulted in heavier biomass and more tubers as had been observed and recorded in this study.

Taller and numerous off-shoots of lowland ecotype *C. rotundus* had very minimal reductions on heights of transplanted and wet direct-seeded rice, corroborating the earlier claim that light was not a factor for competition between weed and rice. Although height was not measured, however, a study showed that leaf transmission ratios and leaf area index of upland rice planted at 0, 60, and 120 kg N ha-^-1^ significantly decreased at increasing population of upland *C. rotundus* (0, 150, 350, 550, and 750 plants m^-2^) [21]. On the other hand, lowland ecotype *C. rotundus* caused greater reductions in the number of leaves, tillers, and panicles of transplanted rice when solar radiation was low (Run II). This suggests that growth adjustment (plasticity) of the weed in response to such condition greatly affects the three growth variables of transplanted rice.

Information on competitive ability helps confirm whether a certain plant is truly a weed of a cultivated plant. It also helps decide whether the execution of control is necessary and economical to be implemented. Our study also demonstrated that lowland ecotype *C. rotundus* could significantly reduce the yield of transplanted and wet direct-seeded rice under flooded conditions. This simply implies that it is a weed not only of upland rice but also of irrigated lowland rice. To our knowledge, this is the first report showing the negative effects of *C. rotundus* on yield of rice under such conditions. Under upland conditions, a study showed that full competition of upland *C. rotundus* reduced the grain yield of drilled and broadcasted rice plants by 42.6 and 51%, respectively [20]. When it competed together with annual weeds, yield of drilled rice was further reduced by 83%; broadcasted rice by 77.6%. In an experiment conducted in the early wet season of 1972, upland *C. rotundus* at 150 to 750 density m^-2^ also reduced the grain yield of upland rice by 15 to 53.8% at 0 kg N ha^-1^, 23 to 44.1% at 60 kg N ha^-1^, and 22 to 47.2% at 120 kg N ha^-1^; in the late wet season of the same year, the weed at 100 to 400 density m^-2^ reduced yield by 7.1 to 21.4% at 0 kg N ha^-1^, 19 to 28.1% at 60 kg N ha^-1^, and 17.5 to 30% at 120 kg N ha^-1^; in the dry season of 1973, the yield was reduced by 6.3 to 25% at 0 kg N ha^-1^, 9.1 to 36.4% at 60 kg N ha^-1^, and 15 to 40% at 120 kg N ha^-1^ [21]

Although statistical analysis showed no significant differences, the negative effects of lowland ecotype *C. rotundus* on grain yields were generally lower on wet direct-seeded rice than on transplanted rice. This suggests that the negative effects of the weed on rice yield could be further reduced through the direct-seeding method. The use of high seeding rates in direct-seeding can also help suppress weed growths and reduce competition through rapid canopy closure [41]. In fact, in a study on the growth response of upland *C. rotundus* to interference by direct-seeding, it was reported that 12 and 24 rice plants pot^-1^ (equivalent to 60 and 120 kg seeds ha^-1^) reduced the weed’s leaf area by 79 and 86%, respectively [38]. Moreover, it also reduced the shoot biomass, tuber production rate, and leaf biomass of the weed. Other studies showing the effects of direct seeding on weeds had been also reported on weedy rice [42]; *Ammania baccifera* L. [43]; *Ludwigia hyssopifolia* (G. Don) Exell. [44]; *Echinochloa glabrescens* Munro ex Hook. f. [45]; and *Amaranthus spinosus* L. as well as *Ludwigia octovalvis* (Jacq.) Raven [46].

The rectangular hyperbolic yield loss model that we used in this study fitted the relationships between growth variables of lowland ecotype *C. rotundus* and grain yield loss of rice [37]. The values of parameter *i* suggest that the first lowland ecotype *C. rotundus* that will grow and compete could cause loss by as much as 0.8% on transplanted rice and 0.6% on wet direct-seeded rice. The values of parameter *a*, on the other hand, suggest that the maximum density, number of off-shoots, and heavier shoot biomass of lowland ecotype *C. rotundus* could cause loss by as much as 75.7% on transplanted rice and 100% on wet direct-seeded rice.

This study confirmed the following: a) lowland ecotype *C. rotundus* is a weed of rice under flooded conditions, b) it can grow and produce numerous off-shoots and tubers when grown with transplanted and wet direct-seeded rice; and c) its growth and reproduction at 22 to 88 tubers m^-2^ can cause significant reductions on growth and grain yields of both transplanted and wet direct-seeded rice under flooded conditions. To know more of its ecology and identify appropriate weed control strategies against it under irrigated lowland conditions, it was recommended that further researches be conducted. For example, a) its growth and reproduction when grown with rice varieties having higher yield and potential competitive qualities (early-maturing, high-tillering, and heavy shoot production); b) its growth and reproduction at different seeding rates of wet direct-seeded rice; and c) its effects on growth and yield at different growth stages of rice (critical period of weed control) are to be studied. Furthermore, values of parameters *a* and *i* via the rectangular hyperbola model must be also evaluated under field conditions to come up with precise predictions on the negative effects of lowland ecotype *C. rotundus* on yields of transplanted and wet direct-seeded rice.

## Acknowledgements

The authors thank Emerson T. Duque and Shelvin Antolin for the support during the conduct of the experiment; Constante T. Briones for the English critique; and the local government units and farmers of La Purisima, Aliaga, Nueva Ecija, Philippines for their help during the collection of tubers in ricefields.

## Supporting information

**S1 Fig. 1**. Grain yield of rice as affected by different densities of lowland ecotype *Cyperus rotundus* under flooded conditions.

